# Conformational Flexibility of p150^Glued^(1-191) Subunit of Dynactin Assembled with Microtubules

**DOI:** 10.1101/623751

**Authors:** C. Guo, J. C. Williams, T. Polenova

## Abstract

Microtubule-associated proteins (MAPs) perform diverse functions in cells. These functions are dependent on their interactions with microtubules. Dynactin, a cofactor of dynein motor, assists the binding of dynein to various organelles and is crucial to the long-distance processivity of dynein-based complexes. The largest subunit of dynactin, the p150^glued^, contains a N-terminus segment that is responsible for the microtubule-binding interactions and long-range processivity of dynactin. We employed solution and magic angle spinning NMR spectroscopy to characterize the structure and dynamics of the p150^glued^ N-terminal region, both free and in complex with polymerized microtubules. This 191-residue region encompasses the CAP-Gly domain, the basic domain and serine-proline-rich (SP-rich) domain. We demonstrate that the basic and SP-rich domains are intrinsically disordered in solution and significantly enhance the binding affinity to microtubules as these regions contain the second microtubule-binding site on the p150^Glued^ subunit. The majority of the basic and SP-rich domains are predicted to be random-coil, while the segments S111–I116, A124–R132 and K144–T146 in the basic domain contain short α-helical or β-sheet structures. These three segments possibly encompass the microtubule binding site. Surprisingly, the protein retains high degree of flexibility upon binding to microtubules except for the regions that are directly involved in the binding interactions with microtubules. This conformational flexibility may be essential for the biological functions of the p150^Glued^ subunit.

**STATEMENT OF SIGNIFICANCE:** Microtubule-associated proteins (MAPs) perform diverse functions in cells. Many of them comprises intrinsically disordered regions, whose structural flexibility are central to microtubule-based cellular functions of MAPs. We employed solution and magic angle spinning NMR spectroscopy to characterize the structure and dynamics of the p150^glued^ N-terminal region encompassing the CAP-Gly domain, the basic domain and serine-proline-rich (SP-rich) domain, both free and in complex with polymerized microtubules. The results reveal that the basic and SP-rich domains are largely unstructured and retains high degree of flexibility upon binding to microtubules except for the regions that are possibly involved in the binding interactions with microtubules. This approach is informative for dynamics studies of intrinsically disordered MAPs and other disordered proteins in large biological assemblies.

## INTRODUCTION

Microtubules (MTs) play fundamental roles in intracellular transport, cell motility, cytoskeletal organization, maintenance of cell shape and separation of chromosomes during mitosis.(1, 2) Microtubule-associated proteins (MAPs) are involved in these cellular activities by interacting with microtubules and/or regulating microtubule dynamics.(3, 4) The dynein motor mediates the retrograde organelle transport along microtubules.(5, 6) Dynactin, a cofactor of dynein, assists binding of dynein to various organelles, especially mRNA, chromosomes, viruses and endomembrane vesicles, and is crucial for the long-distance processivity of dynein-based complexes.(7, 8)

Most of the microtubule-associated proteins have specific microtubule-binding domains that mediate their interactions with MTs. The 1.2 MDa dynactin complex contains 20 subunits (9); the cytoskeleton-associated protein glycine-rich domain (CAP-Gly) at the N-terminus of the largest subunit P150^Glued^ is the microtubule-binding domain of dynactin. Mutations in the CAP-Gly domain lead to severe neurodegenerative diseases, such as Perry syndrome and distal spinal bulbar muscular atrophy (dSBMA).(10) Multiple investigations have been carried out into the structure of the CAP-Gly domain, in order to understand the mechanism of its interaction with microtubules.(11–15)

Other domains of p150^Glued^ subunit are also essential for the dynactin’s function. As reported previously, the basic domain of p150^Glued^ (residues 115-145) has the ability to bind with microtubules in the absence of CAP-Gly domain and to independently skate along MTs in the absence of dynein.(16) In addition, this domain enhances dynein processivity by four fold and is responsible for the long-range motility of dynein and its long-time interactions with microtubules.(16) More specifically, the K-rich domain (132-152) antagonizes the inhibitory effect of the coiled-coil 1 domain (CC1, residues 214-547) in the microtubule-binding affinity of p150^Glued^.(17) In the N-terminal segment of p150^Glued^ (residues 1-144), the N-terminal tail (residues 1-25) and the basic patch (residues 106-144), greatly increase the binding affinity of the segment to microtubules and these patches affect the lateral associations of tubulins by interacting with the tubulin’s E-hook.(13) Despite these studies atomic-resolution information concerning the structure and dynamics of the p150^Glued^ subunit bound to the microtubules is still lacking limiting our understanding of dynactin’s function.

Here, we focus on the N-terminal segment of p150^Glued^ subunit, which includes the CAP-Gly domain, the basic domain and the serine-proline-rich (SP-rich) domain. This functional region of p150^Glued^ comprises the N-terminal residues 1-191 of p150^Glued^ and is dubbed here p150^Glued^(1-191). We performed structure and dynamics investigation of p150^Glued^(1-191) free and in complex with microtubules, by magic angle spinning NMR (MAS NMR) and solution NMR spectroscopy. On the basis of these experiments we determined the likely microtubule binding site and functional domains in the extended region. Our results inform on the conformation and mobility of the basic and the SP-rich domains of p150^Glued^(1-191) free and bound to microtubules, at atomic resolution. The structural and dynamics information of the p150^Glued^(1-191) subunit bound to microtubules gained in this study is beneficial for understanding the structure-function relationships of p150^Glued^ subunit. Our work lays the foundation for the determination of the structure of the dimeric p150^Glued^ subunit bound to polymerized microtubules.

## EXPERIMENTAL

### Materials

Common chemicals were purchased from Fisher Scientific or Sigma-Aldrich. Isotopically labeled chemicals including ^15^NH_4_Cl and U-^13^C_6_-glucose were purchased from Cambridge Isotope Laboratories, Inc. Chromatography columns were purchased from GE Healthcare. Bovine brain tubulin, GTP and Paclitaxel (taxol) were purchased from Cytoskeleton, Inc. 400-mesh copper grids coated with Formval and stabilized with evaporated carbon films were purchased from Electron Microscopy Sciences. The SMT3 fusion vector (pET28b-His_6_-SMT3) and His_6_-Ulp1 protease expression system were generous gifts from Dr. Christopher Lima (Weill Medical College, Cornell University).

### Protein expression and purification

The N-terminal region of p150^Glued^ subunit of *Rattus norvegicus* dynactin that encompasses residues 1-191 was subcloned into the pET28b-His_6_-SMT3 vector using *BamHI* and *XhoI* restriction sites. The construct was transformed into *E.coli* BL21 (DE3) competent cells for expression and purification. A cysteine residue was added to the C-terminal end of p150^Glued^(1-191) for cysteine-based cross-linking. This p150^Glued^(1-191) construct was expressed as His_6_-SMT3-tagged recombinant protein using the above construct. The natural abundance p150^Glued^(1-191) protein was overexpressed in LB media. The *E. coli* cells were grown at 37 °C in LB media after antibiotic selection on kanamycin LB/agar plates. Bacterial cells were induced with 0.4 mM isopropyl-β-1-thiogalactopyranoside (IPTG) at 37 °C for 4 to 5 hours after the OD^600^ value reached 0.6~0.7 and expressed at 37 °C for 4 to 5 hours, and were harvested with centrifugation at 4000 *g* for 30 minutes.

Overexpression of U-^15^N-labeled and U-^13^C,^15^N-labeled p150^Glued^(1-191) was carried out in minimal media as follows. Cells were grown in 2 L of unlabeled LB media at 37 °C and harvested when the OD^600^ reached ~1.4. The cell pellets were pre-washed with M9 minimum media depleted of nitrogen and carbon sources and collected after centrifugation at 4000 *g* for 30 min. The pelleted cells were resuspended in 1 L of M9 media that contained ^15^NH_4_Cl, 2 g U-^13^C_6_ glucose, basal vitamins, 0.1 mM CaCl_2_, and 2 mM MgSO_4_ at cell concentrations fourfold greater than LB media. ^15^NH_4_Cl and 4 g of natural abundance glucose were used for preparation of U-^15^N-labeled p150^Glued^(1-191). After a recovery in labeled M9 media for 1 hour, the cells were induced with 0.8 mM IPTG at 37 °C and expression proceeded for 4 to 5 hours. Cells were harvested with centrifugation at 4000 *g* for 40 min. The pelleted cells were resuspended in phosphate buffered saline (PBS) and stored at −80 °C.

The p150^Glued^(1-191) protein was purified using Ni-affinity chromatography and gel-filtration chromatography. The cells were lysed after addition of DNase, protease inhibitor and 1mM DTT. The supernatant was collected from cell lysis with centrifugation at 15,000 rpm for 30 min and ultracentrifugation at 69,700 g (TLA-120, fixed angle) for 40 min. The protein solution was loaded onto a His-Trap Ni-affinity column (GE Healthcare) and most impurities were removed by a slow gradient to 140 mM of imidazole in 1x PBS, 1 mM DTT, pH 7.4. p150^Glued^(1-191) fused with His_6_-SMT3-tag was eluted at 300 mM to 350 mM of imidazole. The His_6_-SMT3-p150^Glued^(1-191) was digested with ~1 mg SUMO protease (Ulp1) at 4 °C for overnight after the collected fractions were dialyzed against a buffer solution containing 50 mM Tris, 100 mM NaCl, 1mM DTT and 1mM EDTA to an imidazole concentration of lower than 20 mM. The purified p150^Glued^(1-191) was concentrated to the final volume of 2.0 mL (protein concentration of 0.5 mM for U-^13^C,^15^N-p150^Glued^(1-191)) and loaded onto the gel-filtration column (Superdex 75 pg, GE Healthcare). If needed, cation exchange chromatography can be pursued as a third-step purification using Hi-Trap SP HP column (GE Healthcare). After purification, the extended p150^Glued^(1-191) protein solution was exchanged into the phosphate buffer and concentrated to a final concentration of 0.4 mM for preparation of complexes with MTs. The typical yield of pure p150^Glued^(1-191) was 25 mg per 1 L of LB for unlabeled protein, 20 mg per 1 L M9 media for U-^15^N-p150^Glued^(1-191) and 13 mg per 1 L M9 for U-^13^C, ^15^N-p150^Glued^(1-191).

### Preparation of p150^Glued^(1-191)/MT complexes

Paclitaxel-stabilized microtubules were prepared by polymerization of lyophilized bovine tubulin (Cytoskeleton, Inc). Precleared tubulin was dissolved in 80 mM PIPES, 1 mM MgCl_2_, 1mM EGTA, pH 6.8 (BRB 80). Tubulin was typically incubated at 37 °C for 30 mins with 20 μM paclitaxel, 1 mM GTP, 10 % DMSO in BRB 80 buffer. Polymerized microtubules were pelleted by ultracentrifugation at 108,900 *g* (50,000 RPM) for 20 minutes and resuspended in 20 mM phosphate buffer containing 50 mM NaCl and 1 mM DTT, pH 6.0 for cosedimentation assays and TEM characterization.

To measure the binding affinity of p150^Glued^(1-191) to microtubules, the co-sedimentation assay was performed at molar ratios of p150^Glued^(1-191) to tubulin dimer ranging from 1:1 to 8:1. The concentration of microtubules was typically 20 μM and concentrations of p150^Glued^(1-191) were varied from 0 μM (control) to 160 μM. After the mixture was incubated at 25 °C for 1 hour, the p150^Glued^(1-191)/MT complex was pelleted with ultracentrifugation at 108,900 *g* for 20 minutes. The contents of supernatant and pellets were characterized by SDS-PAGE analysis using 15% acrylamide gel.

For solution NMR experiments, the p150^Glued^(1-191) sample with a final protein concentration of 250 μM was prepared in 500 μL phosphate buffer (20 mM phosphate, 50 mM NaCl, 1 mM DTT and 10 % D_2_O, pH 6.0).

For solid-state NMR samples of p150^Glued^(1-191)/MT assemblies, polymerized microtubules were prepared from 6 mg of lyophilized tubulin. The resulting solution was directly resuspended in the p150^Glued^(1-191) solution at a 1:2 molar ratio of tubulin dimer to p150^Glued^(1-191) and containing 10% sucrose cushion. After 1 hour incubation at 25 °C, the assemblies were pelleted by ultracentrifugation at 108,900 *g* and transferred into an MAS rotor. 9.6 mg hydrated p150^Glued^ (1-191)/MT complex was packed in a 1.6 mm Varian rotor.

For ^31^P MAS NMR experiments, 8.5 mg polymerized microtubules and 6.3 mg CAP-Gly^19-107^/MT complex containing ~1 mg U-^13^C,^15^N-CAP-Gly^19-107^ were packed in two 1.6 mm Varian rotors. Both samples were hydrated. To avoid any background ^31^P signals from buffers (^31^P is a 100% natural abundant nucleus), the CAP-Gly^19-107^ solution was exchanged into BRB15 buffer (15 mM PIPES, 1 mM MgCl_2_, 1 mM EGTA, pH 6.8) and the CAP-Gly^19-107^/MT complex was prepared in BRB15 buffer instead of phosphate buffer. Sodium phosphate dibasic heptahydrate (Na_2_HPO_4_·7H_2_O) was packed in a 1.6 mm Varian rotor as a standard compound.

### Transmission electron microscopy

The morphologies of polymerized microtubules and their complexes with p150^Glued^(1-191) and CAP-Gly^19-107^ were characterized by negatively stained TEM with a final concentration of microtubules of 5-10 μM. The microtubule samples were stained with uranyl acetate (5% w/v) and deposited onto 400 mesh copper grids coated with formval and carbon films. The TEM images were acquired by a Zeiss Libra 120 transmission electron microscope operating at 120 kV.

### Solution NMR spectroscopy

All solution NMR spectra were acquired on a 14.1 T (^1^H Larmor frequency of 600.1 MHz) Bruker Avance spectrometer using a triple-resonance (HCN) CryoProbe at 298 K. The 2D HSQC spectra were recorded with U-^15^N-p150^Glued^(1-191) and the 3D spectra were acquired with U-^13^C, ^15^N-p150^Glued^(1-191). For the H-N HSQC experiment, 256 complex points were recorded in the ^15^N dimension; States-TPPI was used for phase sensitive detection. The 3D HNCA and HN(CO)CA spectra were acquired in the States-TPPI mode(18) in both indirect dimensions and ^15^N decoupling was applied during the acquisition. For the HNCA experiment, 104 complex points were recorded in indirect ^13^C and ^15^N dimensions. For the HN(CO)CA experiment, 112 points and 100 points were recorded in the ^13^C and ^15^N dimension, respectively. A recycle delay of 1.0 s was used, and the experimental time was 2 days for both 3D spectra. The WATERGATE sequence(19) was applied to suppress the ^1^H signals arising from water. Chemical shifts in all spectra were referenced to DSS. All NMR spectra were processed using NMRPipe(20) and analyzed in CCPNMR.(21)

### MAS NMR spectroscopy

^**31**^P and ^13^C-detected MAS NMR spectra were collected at 14.1 T on a narrow bore Varian InfinityPlus instrument using a 1.6 mm triple resonance HXY probe. The Larmor frequencies were 599.5 MHz for ^1^H, 242.8 MHz for ^**31**^P, 150.7 MHz for ^13^C and 60.7 MHz for ^15^N. The MAS frequency was 14 kHz controlled at ± 10 Hz for all experiments using the Varian MAS controller. The sample temperature was calibrated using KBr as the temperature sensor.(22)

For ^31^P experiments, the sample temperature was approximately 5.6 °C for paclitaxel-stabilized MTs, 7.1 °C for CAP-Gly^19-107^/MT complex, and 26.0 °C for Na_2_HPO_4_·7H_2_O, maintained to within ± 0.2 °C by a Varian temperature controller. The typical 90° pulse lengths were 2.5 μs for ^31^P and 2.4 μs for ^1^H. The Na_2_HPO_4_·7H_2_O was used as the standard to calibrate the pulse lengths as well as to optimize the ^1^H–^31^P CP Hartmann-Hahn matching conditions. The ^31^P direct-excitation experiments were performed with recycle delays ranging from 10 to 15 s, to assure the equilibrium magnetization recovery. For ^1^H-^31^P CPMAS experiments, the CP contact time was 2.5 ms. TPPM(23) with radio frequency (rf) field strength of 100-110 kHz was used for decoupling during all experiments. The ^31^P chemical shift was referenced to H_3_PO_4_ (85% H_3_PO_4_ in H_2_O), used as an external referencing standard.(24)

In ^13^C-detected MAS NMR experiments on p150^Glued^(1-191)/MT assemblies a series of 1D ^1^H–^13^C CPMAS spectra were acquired at the sample temperatures of −27.0 °C −25.8 °C, −24.5 °C, −17 °C, −10 °C and 4 °C. The typical 90° pulse lengths were 2.5 μs for ^1^H, 3.5 μs for ^13^C and 3.0 μs for ^15^N. During the ^1^H–^13^C CP transfer, a linear amplitude ramp of 80–100% was applied on the ^1^H channel. The RF field strength was ~100 kHz for ^1^H TPPM decoupling during the acquisition period. The 2D ^13^C-^13^C correlation spectra were acquired at several sample temperatures ranging from −27 °C to 4 °C. TPPI(25) acquisition mode was used for phase sensitive detection in the indirect dimension of all 2D spectra. In the CORD(26) experiments, the ^1^H rf fields during the ^13^C-^13^C mixing were matched to the spinning frequency (14 kHz) or half of the spinning frequency (7 kHz) and the mixing time was 50 ms. All 2D CORD spectra were processed with either 90° or 60° shifted sine-bell function in both dimensions.

The 2D heteronuclear correlation NCA and NCO spectra were acquired at a sample temperature of −24.5 °C. The ^1^H-^15^N CP was performed with the rf fields of 97 kHz on ^1^H and 83 kHz on ^15^N, respectively; a linear amplitude ramp of 80-100% was applied on ^1^H. For SPECIFIC CP, the Hartmann-Hahn condition was matched at a ^13^C rf field strength of 35 kHz and a ^15^N rf field strength of 63 kHz using a tangent amplitude ramp. The carrier frequencies in the ^13^C dimension were set to 55 and 170 ppm in the NCA and NCO experiment, respectively.

## RESULTS AND DISCUSSION

### Biochemical Characterization of p150^Glued^(1-191) Binding with Polymerized Microtubules

The p150^Glued^(1-191) was isolated as a monomer in a two-step purification procedure, as shown in Fig. 1B and Fig. S1A. The binding affinity of monomeric p150^Glued^(1-191) to polymerized microtubules was assessed by co-sedimentation assay (Fig. 1C). More than 98% of p150^Glued^(1-191) binds to microtubules when equivalent molar concentrations of the protein and the tubulin dimer were used (Fig. 1C). In comparison, as we and others reported previously, under the identical conditions less than half of the CAP-Gly domain protein encompassing residues 19-107 (CAP-Gly^19-107^) binds to MTs.(15) Thus the binding affinity of p150^Glued^(1-191) to MTs is significantly higher than that of CAP-Gly^19-107^. This result validates that p150^Glued^(1-191) contains a second MT-binding domain other than CAP-Gly and the extended region comprised of the basic domain and SP-rich domain assists the interactions between the p150^Glued^ and MTs.(16) Furthermore, the amount of p150^Glued^(1-191) bound to MTs increases as molar ratio of p150^Glued^(1-191) to tubulin dimer becomes higher, indicating that the binding interaction between p150^Glued^(1-191) and MTs is promoted by the higher molar concentration of the protein. The fraction of MT-bound p150^Glued^(1-191) decreases when the molecular ratio of p150^Glued^(1-191) to tubulin dimer is higher than 2:1. Therefore, the p150^Glued^(1-191)/MT assemblies were prepared with the 2:1 p150^Glued^(1-191): tubulin dimer molar ratio, to ensure the stoichiometric formation of the assembly.

**Figure 1.**
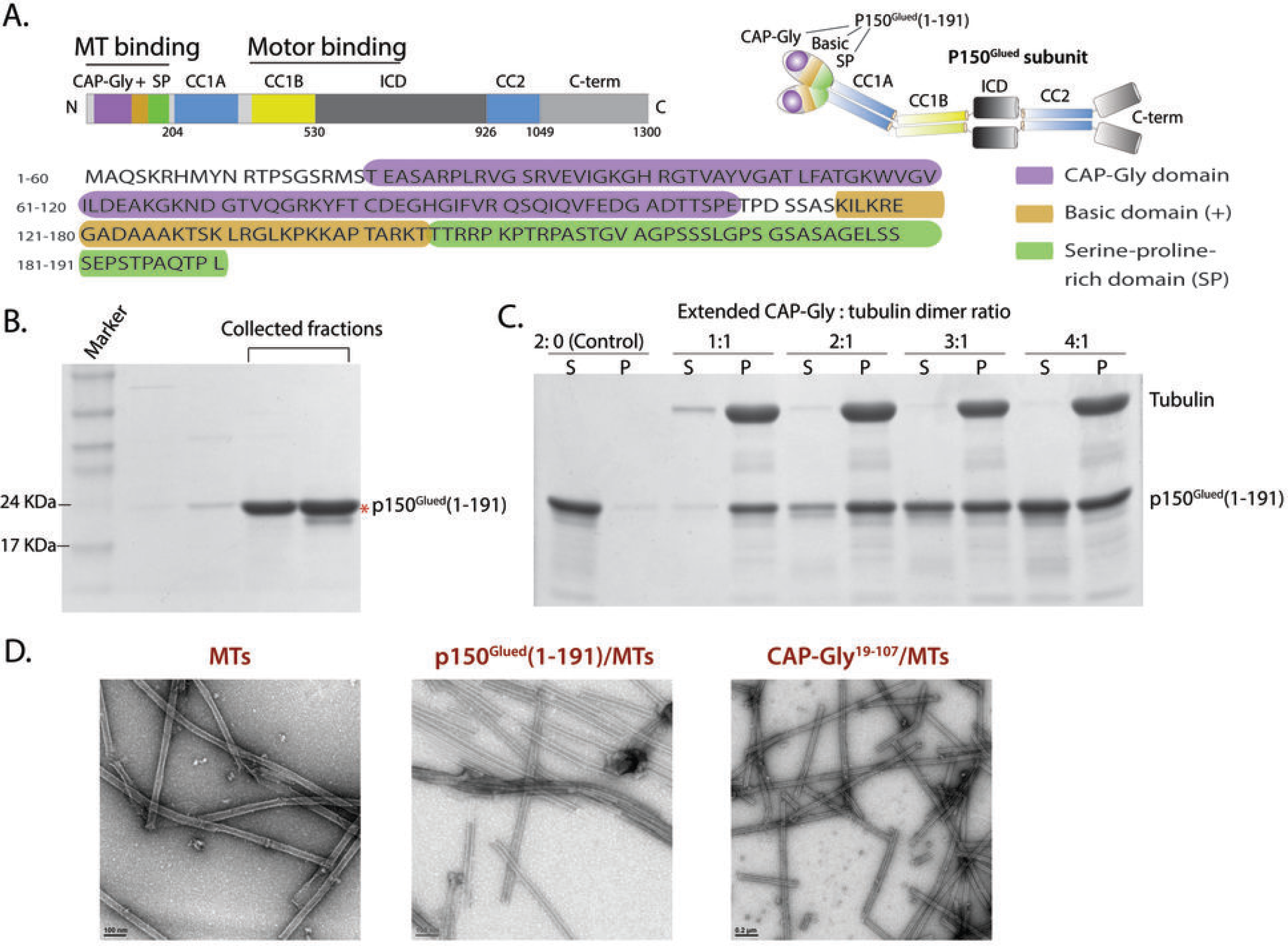
A. *(Top)* Domain structure of p150^Glued^ subunit of dynactin. (Bottom) Primary sequence of p150^Glued^(1-191) domain. The p150^Glued^(1-191) consists of the CAP-Gly domain (19-107), basic domain (115-145) and serine-proline-rich domain (146-191). The basic and SP-rich domains are located in the linker connecting the CAP-Gly and coiled-coil 1 domains (CC1). B. SDS-PAGE gel of p150^Glued^(1-191) after purification. C. Co-sedimentation assay of p150^Glued^(1-191) with MTs. “S” and “P” denote supernatant and pellets after ultracentrifugation of the p150^Glued^(1-191)/MT solution, respectively. D. Negatively stained TEM images of MT alone (left), p150^Glued^(1-191)/MT assemblies (middle) and CAP-Gly/MT assemblies (right).

The p150^Glued^(1-191) was also isolated as a cross-linked dimer by dialysis against a buffer solution containing no DTT, see Fig. S1B of the Supporting Information. The dimer appears to bind to polymerized MTs with higher binding affinity than the monomer (Fig. S1C). While in this work we have not pursued structural characterization of the dimeric p150^Glued^(1-191) complex with microtubules, such studies will be conducted in the future.

To assess the morphology of the assembly of p150^Glued^(1-191) bound with polymerized microtubules, we used negatively stained TEM. As shown in Fig. 1D, p150^Glued^(1-191) bound to polymerized microtubules forms straight tubules, of the same dimensions (within the resolution of the images) and morphology as the pure microtubule preparations as well as the assembly of CAP-Gly(19-107) with polymerized microtubules.

### ^31^P MAS NMR spectroscopy of microtubules and CAP-Gly^19-107^/microtubule assembly

We have recorded ^31^P MAS NMR spectra to characterize the nucleotide states in microtubules and assess the conformational homogeneity of the microtubules in the samples under investigation. As shown in Fig. 2C, the spectra of microtubules are well resolved, and signals from the GTP and GDP are present and well resolved. Specifically, the direct-excitation and/or CPMAS spectra contain three resonances corresponding to GTP, two corresponding to GDP and two corresponding to the inorganic phosphate PO_4_^−^ (Pi). The signal assignments are shown in Fig. 2C and Table S1 of the Supporting Information. The intensities of the Pα and Pβ resonances of GTP are higher than those of the Pα’ and Pβ’ resonances of GDP. This is consistent with the fact that there are more GTP than GDP molecules bound. In polymerized microtubules, the α-tubulin binds to GTP and β-tubulin binds to either GTP or GDP with a “GTP cap” at the plus-end of MTs that contains the GTP-bound β-tubulins. The two resonances of Pi indicate the potential existence of stabilized GDP-Pi-capped microtubules in the sample of free microtubules.(27–29) Important to note is that the ^31^P peaks in all spectra are narrow (the line widths are 0.4-0.8 ppm) indicating that the conformational homogeneity of the microtubules in our preparations is high, both in the free and CAP-Gly^19-107^ bound states.

**Figure 2.**
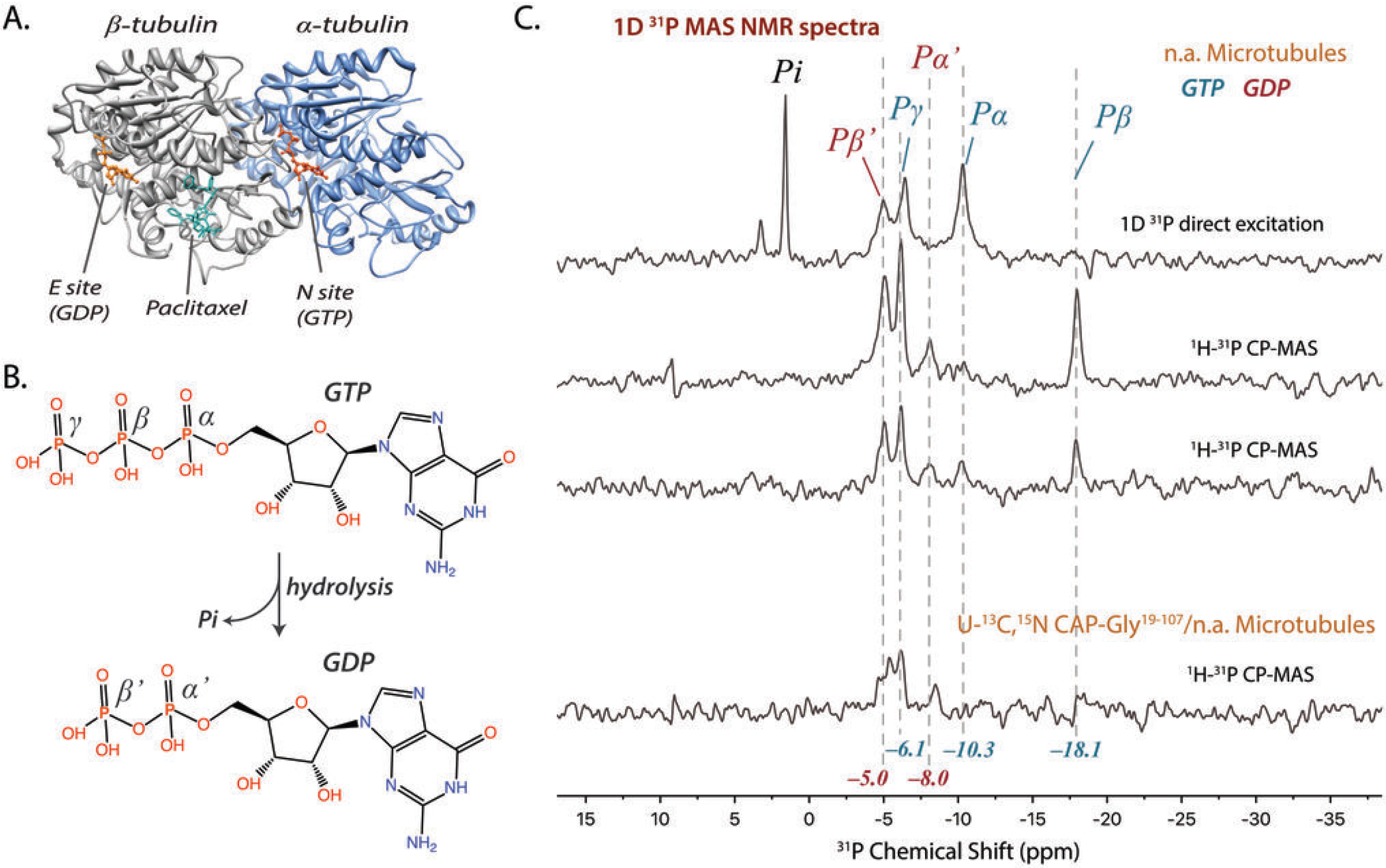
A. The structure of a tubulin heterodimer, showing the positions of GTP, GDP and paclitaxel (PDB ID: 1JFF(34)). B. Chemical structures of GTP and GDP, indicating the phosphorus atoms in each molecule. C. ^31^P MAS NMR spectra of guanosine nucleotides in microtubules and CAP-Gly/MT complex. The direct-excitation and CPMAS spectra contain three resonances corresponding to GTP, two corresponding to GDP and two corresponding to the inorganic phosphate PO_4_^−^ (Pi). Each peak was assigned to corresponding phosphorus resonance of GTP and GDP. The ^31^P MAS NMR spectra were acquired with (from top to bottom) 20,000 scans, 4864 scans, 8360 scans and 7756 scans.

### Solution NMR spectroscopy of p150^Glued^(1-191)

The 2D ^1^H-^15^N HSQC and 3D HNCA and HN(CO)CA spectra of U-^15^N-p150^Glued^(1-191) and U-^13^C,^15^N-p150^Glued^(1-191) are shown in Fig. 3. The ^1^H and ^15^N chemical shift perturbations with respect to the free CAP-Gly domain are summarized in Table S3. In the 2D HSQC spectra, the signals that correspond to residues in the CAP-Gly domain of p150^Glued^(1-191) are well dispersed and generally overlay well with those in the spectra of CAP-Gly^19-107^ reported by us previously,(15) see Fig. 3A. The residues with large ^15^N chemical shift perturbations (CSPs) are S19, E21, A22 (CSPs greater than 0.4 ppm) as well as T20, S23, A24, Q95-F97, T104, S105 (CSPs greater than 0.2 ppm but smaller than 0.4 ppm) (Fig. 4A). All of these residues are located at the N- and C-termini of CAP-Gly domain (Fig. 4 C), so these changes are not surprising. These residues are more rigid and structured in the context of the p150^Glued^(1-191) protein compared to the CAP-Gly domain alone. In addition to these perturbations, several residues in the β-sheet core region of CAP-Gly exhibit small chemical shift changes commensurate with minor structural rearrangements; these are R25, R28, R41, N69, H85 and R90, see Fig. 4C, which are mostly positively charged residues.

**Figure 3.**
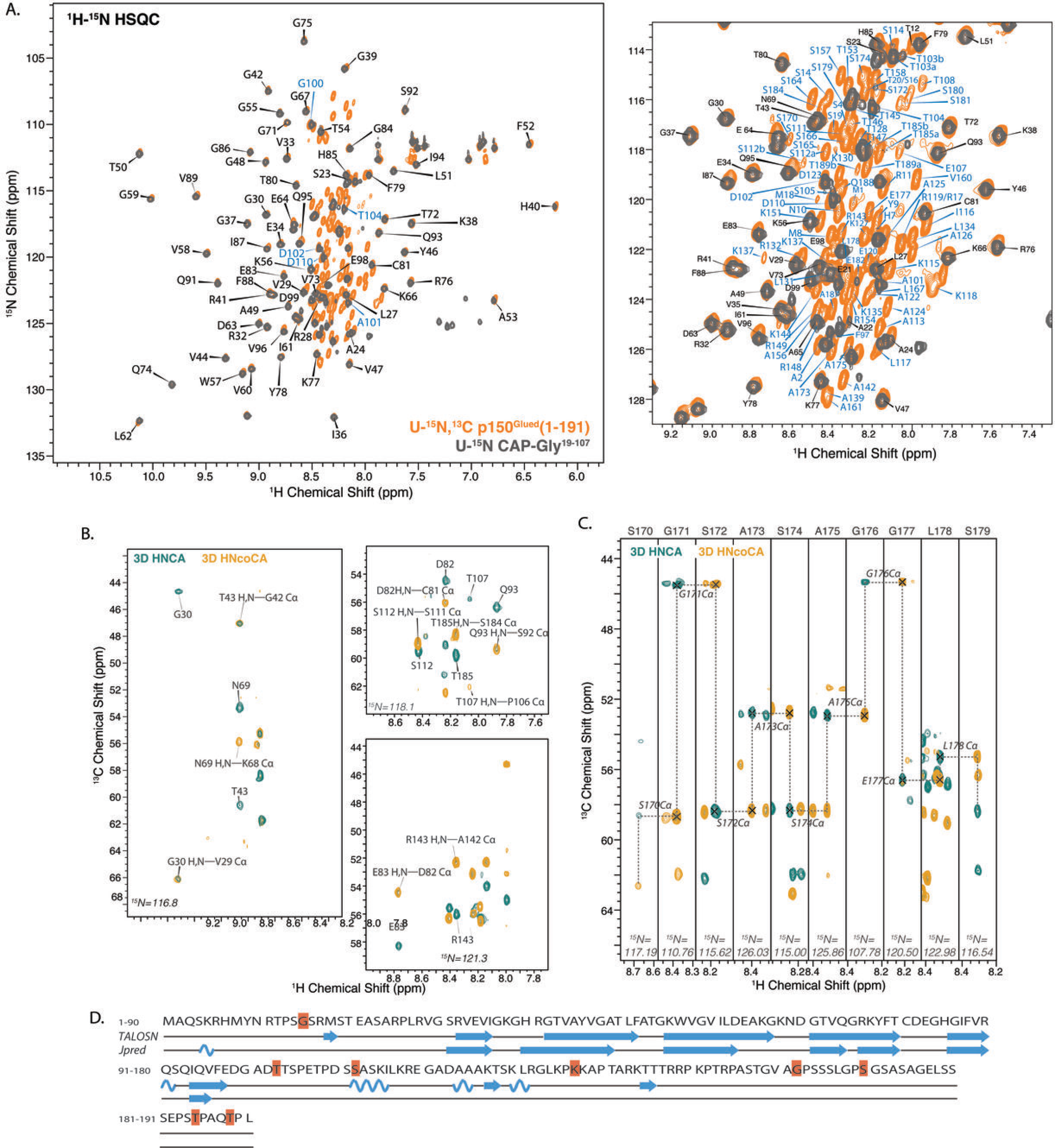
Solution NMR spectra of p150^Glued^(1-191). A. (Left) Superposition of 2D ^1^H-^15^N HSQC spectra of U-^13^C,^15^N-p150^Glued^(1-191) (orange) and U-^13^C,^15^N CAP-Gly^19-107^ (gray). (Right) An expanded view of 2D HSQC spectra showing the resonance assignments of the poorly dispersed regions including the basic and SP-rich domains. Residues T103, S112, T185 and T189 show double peaks. The assignments of cross peaks corresponding to residues in different domains are color coded as follows: CAP-Gly domain, black; extended regions including basic domain and serine-proline-rich domain, blue. B. Representative 2D ^1^H-^15^N planes of 3D HNCA (cyan) and HNcoCA (yellow) solution NMR spectra. C. Sequential backbone walk for residues S170–S179 using the 3D solution NMR spectra. D. Secondary structure of p150^Glued^(1-191) predicted by TALOS-N based on solution NMR chemical shifts compared to that predicted by Jpred from the primary sequence.

In contrast to the CAP-Gly domain, the signals of other domains in the protein are poorly dispersed. This is an indication that the extended region of p150^Glued^(1-191) comprising the basic and SP-rich domains is much less structured than the CAP-Gly domain and may have a substantial proportion of random coil. We have assigned the resonances on the basis of the 3D HNCA and HN(CO)CA spectra. Both intra-residue and sequential correlations are revealed in the HNCA spectra. As shown in Fig. 3B-C, the spectral resolution is excellent permitting full assignments, despite the presence of many Pro residues and sequence repeats. The solution chemical shits are summarized in Table S2 of the Supporting Information.

**Figure 4.**
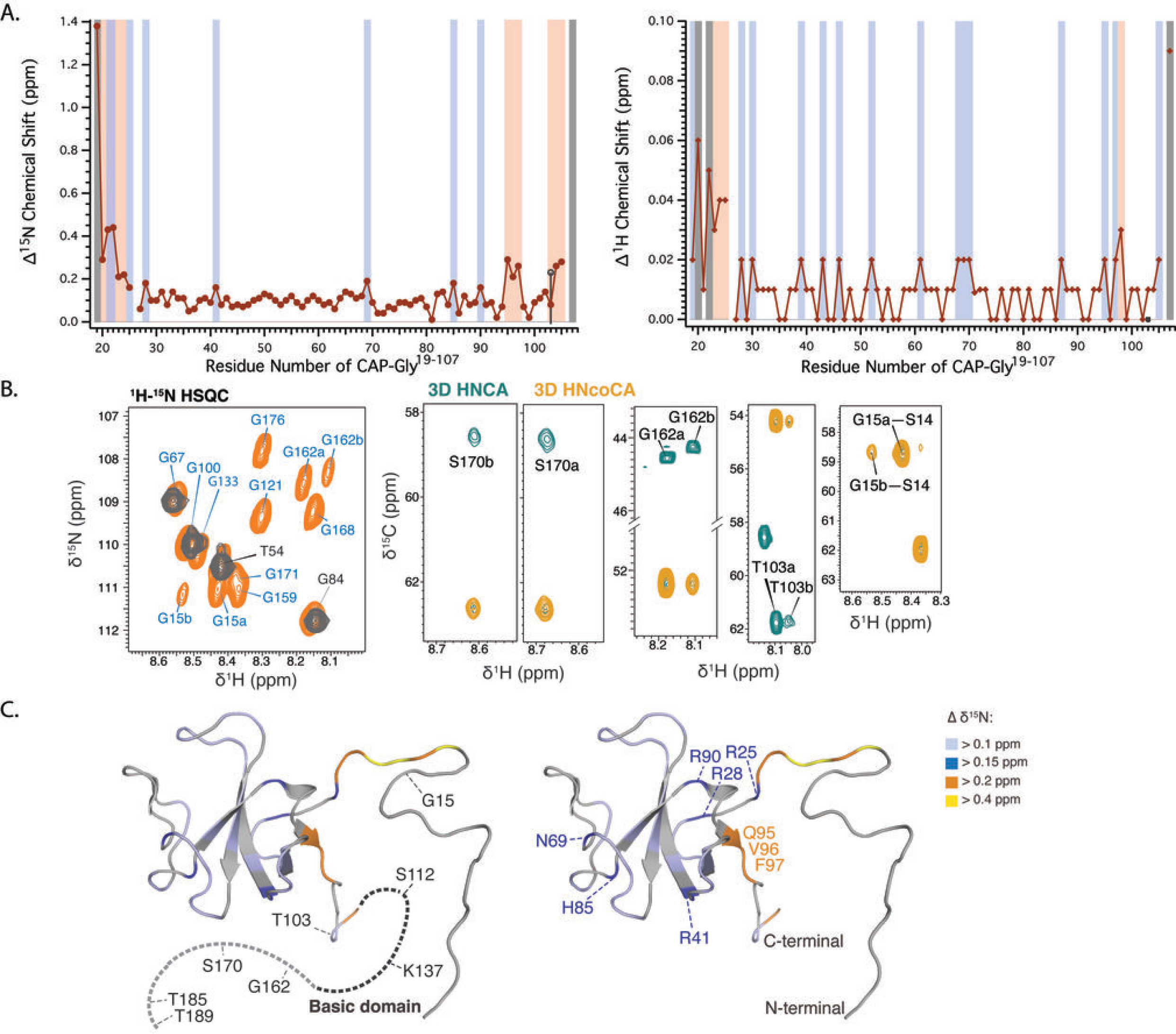
A. Chemical shift perturbations of the CAP-Gly domain plotted against residue number in p150^Glued^(1-191) vs. CAP-Gly^19-107^. The residues are color coded as follows: residues with ^15^N chemical shifts perturbation (∆^15^N) larger than 0.15 ppm or with ^1^H chemical shifts perturbation (∆^1^H) larger than 0.02 ppm, light blue; residues with ∆^15^N larger than 0.2 ppm or with ∆^1^H larger than 0.03 ppm, orange; residues with ∆^15^N larger than 0.4 ppm, light purple; residues with ∆^15^N larger than 1.0 ppm or with ∆^1^H larger than 0.05 ppm, gray. B. Glycine regions of 2D HSQC and 2D slices of 3D spectra showing peak doubling for representative residues in p150^Glued^(1-191). Residues that exhibit two resonances include G15, T103, S112, K137, G162, S170, T185 and T189 (marked in red in Figure 3). C. Mapping of residues exhibiting two resonances *(left)* and ^15^N chemical shift perturbations *(right)* onto 3D CAP-Gly structure (PDB ID:2COY(35)). The residues are color coded as follows: residues with ^15^N chemical shifts perturbation (∆^15^N) larger than 0.4 ppm, yellow; residues with ∆^15^N larger than 0.2 ppm, orange; residues with ∆^15^N larger than 0.15 ppm, dark blue; residues with ∆^15^N larger than 0.1 ppm, light purple.

We have carried out the secondary structure prediction on the basis of the chemical shifts, using TALOS-N(30) and Jpred(31) programs. As shown in Fig. 3D, the predictions for the core regions in the CAP-Gly domain are consistent with the secondary structure of isolated CAP-Gly^19-107^, which is comprised of loop regions and a core region formed by four β-sheets. The SP-rich domain and the majority of the basic domain are random coil. The exceptions are three separate segments in the basic domain, S111–I116, A124–R132 and K144–T146, which are predicted to contain short α-helical or β-sheet stretches.

It is worth noting that several residues exhibit peak doubling in solution NMR spectra (Fig. 4 B and right panel of Fig. 3A), indicating the existence of conformational isoforms for the segments that contain these residues. These residues include G15, T103, S112, K137, G162, S170, T185 and T189, which are mostly in the random coil or loop regions.

Our results thus indicate that despite of the presence of some secondary structures in these segments, the basic and SP-rich domains are intrinsically disordered and unfolded in free state in solution.

### MAS NMR spectroscopy of p150^Glued^(1-191) in complex with microtubules

As shown in Fig. 5A, 1D ^1^H-^13^C CPMAS spectra of U-^13^C,^15^N-p150^Glued^(1-191) assembled with polymerized microtubules, show strong temperature dependence, indicating the presence of dynamics. The spectral intensity and resolution depend on whether a particular temperature was reached by heating or cooling the sample, in the range of 4 °C to −27 °C, which is consistent with our previous observations in CAP-Gly^19-107^ free and assembled with MTs as well as LC8.(11, 12, 14, 32) Different spectra were observed upon temperature cycling, as shown in Fig. 5A. Specifically, at the starting temperature of 4 °C spectral resolution is excellent but deteriorates progressively accompanied by an increase in sensitivity as the experimental temperature is decreased. At −27 °C, the signal-to-noise ratio of the aliphatic and carbonyl ^13^C signals increases by 4- and 8- fold, respectively, compared to that at 4 °C. When the temperature is raised from −25.8 °C to −10 °C after the sample was cooled down to −27 °C, the resolution and sensitivity of the ^1^H-^13^C CPMAS spectra do not change dramatically. The increase in resolution is only observed at −25.8 °C when warming up the sample from −27 °C.

**Figure 5.**
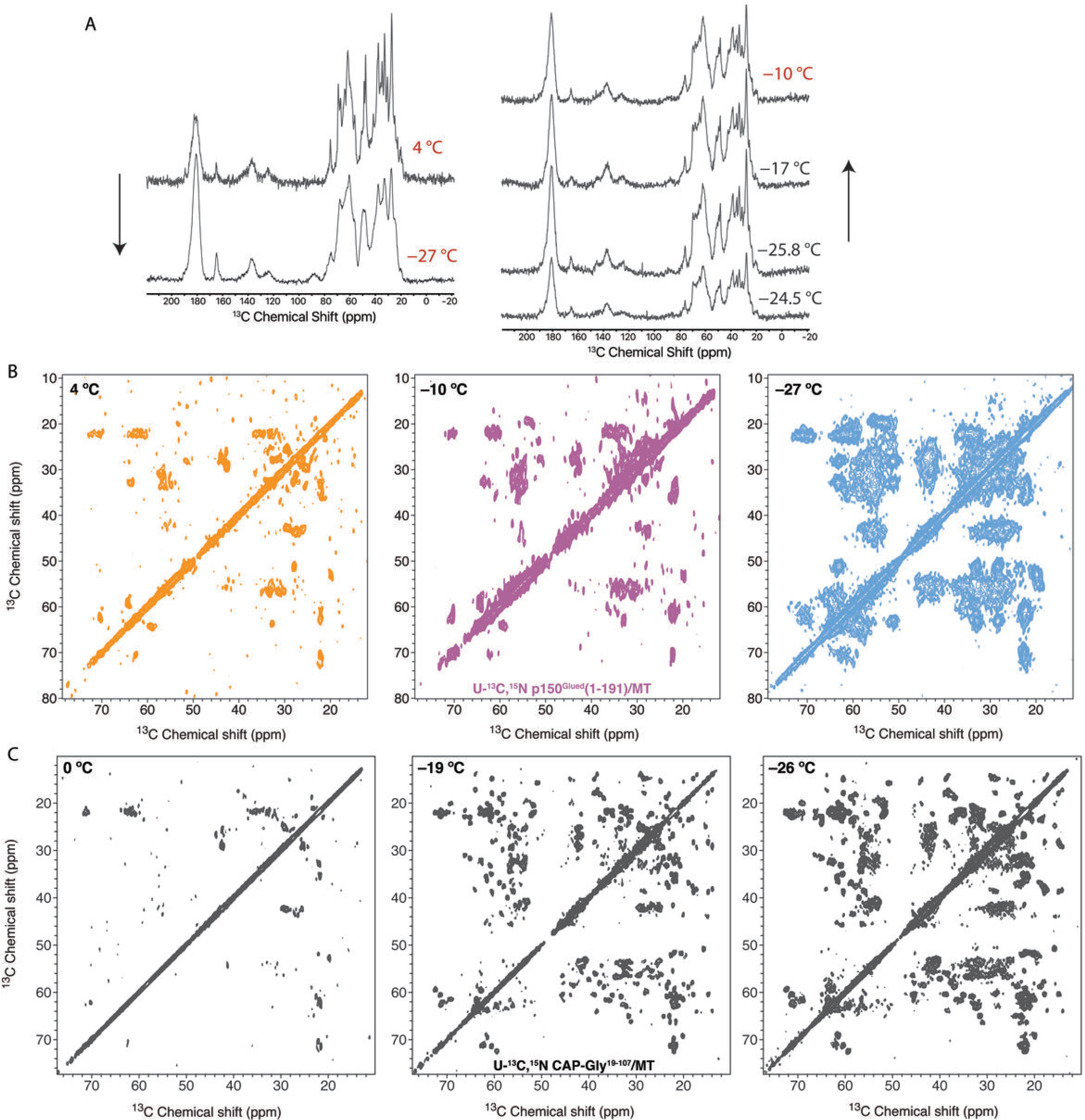
A. Temperature dependence of 1D ^1^H-^13^C CPMAS spectra of U-^13^C,^15^N-p150^Glued^(1-191)/MT complex. *(left)* Spectral resolution is excellent at 4 °C but deteriorates progressively accompanied by increase in sensitivity as the experimental temperature is decreased to −27 °C. The number of transients is 512 for +4 °C and 128 for −27 °C. *(right)* The spectral resolution and sensitivity does not change dramatically when the temperature is raised from −25.8 °C to −10 °C after the sample has been cooled down to −27 °C. The number of transients is 128 for −24.5 °C and 256 for −25.8 °C, −17 °C and −10 °C. B,C. Two dimensional ^13^C-^13^C correlation MAS NMR spectra of U-^13^C,^15^N-p150^Glued^(1-191)/microtubules (B) and U-^13^C,^15^N-CAP-Gly^19-107^/MT (C) at different temperatures. At near or above 0 °C, a finite number of correlations are detected for both p150^Glued^(1-191)/MT and CAP-Gly^19-107^/MT due to motions of flexible regions. For p150^Glued^(1-191)/MT complex, a large number of correlations only appear at −27 °C since motions of residues in flexible regions are restricted. The contour levels were set to 3x noise level. All 2D ^13^C-^13^C spectra were processed with 60° shifted sine-bell function.

Interestingly, the temperature profile of the p150^Glued^(1-191)/MT complex is different from that for the CAP-Gly^19-107^/MT complex, indicating different dynamic behavior. Specifically, the CPMAS spectra of the CAP-Gly^19-107^/MT complex at temperatures close to −10 °C experience significant loss of sensitivity and resolution, while the sensitivity and resolution in the CPMAS spectra of extended p150^Glued^(1-191)/MT complex at −10 °C are the same as at −25.8 °C. This result suggests overall reduced dynamics in the p150^Glued^(1-191) in complex with MTs compared to that in the CAP-Gly^19-107^/MT complex, on timescales of micro- to milliseconds throughout the entire protein molecule.

To further probe the relative flexibility of different segments in p150^Glued^(1-191) assembled with MTs, 2D ^13^C-^13^C CORD spectra were acquired at −27 °C, −10 °C and 4 °C (Fig. 5 B). Although the spectrum acquired at 4 °C exhibits good resolution, only a finite number of correlations are present, due to motions of flexible regions occurring at temperatures above 0 °C. This behavior is similar to that of CAP-Gly^19-107^ and indicates that the limited number of residues detected at 4 °C belong to rigid regions of p150^Glued^(1-191) assembled on microtubules. At −27 °C, numerous intense cross peaks are present in the spectra, consistent with efficient dipolar-based transfers and spin diffusion, which indicates that the motions are restricted under these conditions. The majority of these correlations are only detected at −27 °C and disappear in the ^13^C-^13^C spectra at −10 °C, providing evidence that these are associated with residues in the flexible segments of the protein.

We next pursued resonance assignments of the p150^Glued^(1-191)/MT complex, on the basis of the 2D homo- and heteronuclear correlation spectra, shown in Fig. 6. For this, we used solution NMR chemical shifts as guides.(33) We first assigned the unique signals in 2D heteronuclear NCA spectra, which comprise ~37 % of residues, of which half are located at CAP-Gly domain and half- in other regions. Next, we analyzed 2D ^13^C-^13^C spectra, on the basis of which we made assignments for additional 68 residues. Most of the signals in the CORD spectra acquired at 4 °C were unambiguously assigned except for several signals of high intensity, which possibly contain multiple peaks. Comparison of CORD spectra acquired at −27 °C and −10 °C was very useful for distinguishing residues in the flexible segments of the protein, as shown in Fig. 6C. Specifically, the correlations detected only at −27 °C correspond to the flexible regions of the protein (Fig. 6E), while those at −10 °C - to more rigid residues. Correlations only present at 4 °C correspond to residues in the most rigid segments (Fig. 6B). As shown in Fig. 6E, the flexibilities of different regions in p150^Glued^(1-191) are found to be generally consistent with the predicted secondary structure by TALOS-N. In the basic domain, three segments are predicted to have α-helical or β-sheet structures, S112–R119, A126–R312 and T141–T146. Most of these residues are present in the spectra at 4 °C. At the same time, the SP-rich domain is comprised of flexible segments, both according to the secondary structure predictions and the presence/absence of cross peaks in the CORD spectra acquired at different temperatures.

**Figure 6.**
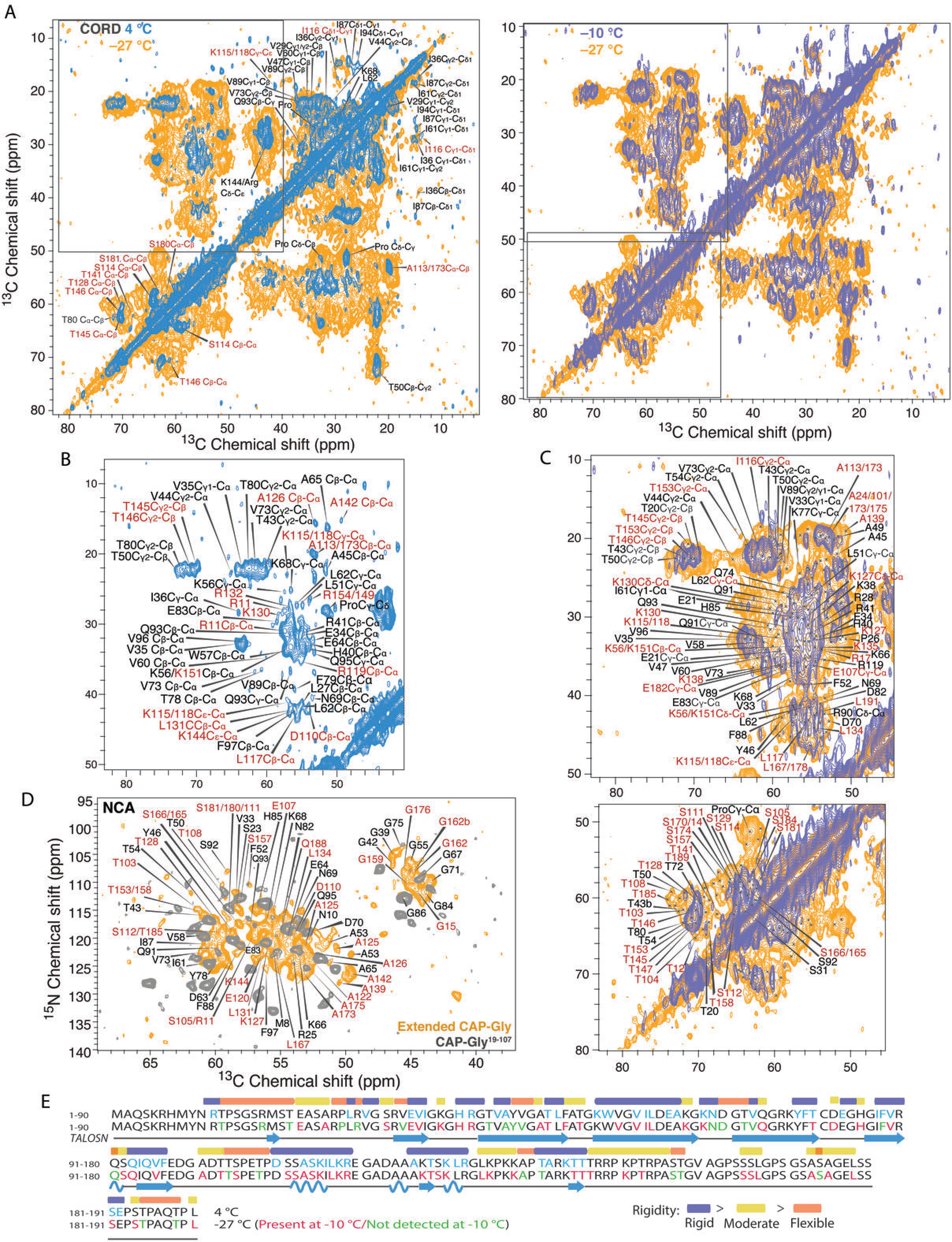
MAS NMR spectra of U-^13^C,^15^N-p150^Glued^(1-191)/microtubules. A. *(left)* 2D ^13^C-^13^C CORD spectrum of U-^13^C,^15^N-p150^Glued^(1-191)/MTs acquired at 4 °C (blue) superimposed with the CORD spectrum acquired at −27 °C (orange). *(right)* 2D CORD spectrum acquired at −10 °C (purple) superimposed with the CORD spectrum acquired at −27 °C. The assignments of cross peaks corresponding to residues in different domains are color coded as follows: CAP-Gly domain, black; extended regions including basic domain and serine-proline-rich domain, red. B. Expanded view of the 2D ^13^C-^13^C CORD spectrum acquired at 4 °C. C. 2D ^13^C-^13^C CORD spectrum acquired at −27 °C showing resonance assignments. The 2D CORD spectrum recorded at −10 °C was superimposed to show the correlations that are present at −10 °C. D. 2D ^15^N-^13^C NCA correlation MAS NMR spectra of U-^13^C,^15^N-p150^Glued^(1-191)/MT (orange) and U-^13^C,^15^N-CAP-Gly^19-107^/MT (gray). E. Primary sequence of p150^Glued^(1-191) showing the relative flexibilities of the different regions of the protein. The correlations detected only at −27 °C correspond to the flexible segments. Resonances detected only at −10 °C are associated with moderately flexible residues, and those present at 4 °C indicate rigid residues. The color coding of the individual residues is as follows: residues appearing at 4 °C, cyan; residues appearing only at −27 °C, green; residues appearing at both −10 °C and −27 °C, pink. The color coding of the segments of secondary structure is as follows: rigid, purple; flexible, light red; moderately flexible, yellow.

Our results thus indicate that the rigid segments in the basic domain of p150^Glued^(1-191), S112–R119, A126–R312 and T141–T146, likely comprise the intermolecular interface with microtubules. (Fig. 7) Together with the CAP-Gly domain, they appear to be responsible for the MT-binding interactions of the p150^Glued^ subunit. This finding is consistent with previous studies demonstrating that the basic domain is the second MT-binding segment of p150^Glued^.(16) Furthermore, the K-rich segment R132-P152 represses the inhibitory effect of the CC1 domain on the MT binding, whereas the SP-domain and CC1 do not bind to MTs.(17) We also speculate that the N-terminal tail (1-25) of p150^Glued^(1-191) may contain an anchoring point at residue R11 for interacting with the tubulin’s C-terminal tail (Fig. 7). In a previous study, the N-terminal segment 1-25 has been proposed to secure the binding of CAP-Gly to microtubules by wrapping around tubulin’s E-hook.(13) According to our results, R11 is the only residue that is detected at 4 °C and exhibits strong peak intensity in the segment 1-25, indicating that R11 is possibly involved in binding interactions with tubulin.

**Figure 7.**
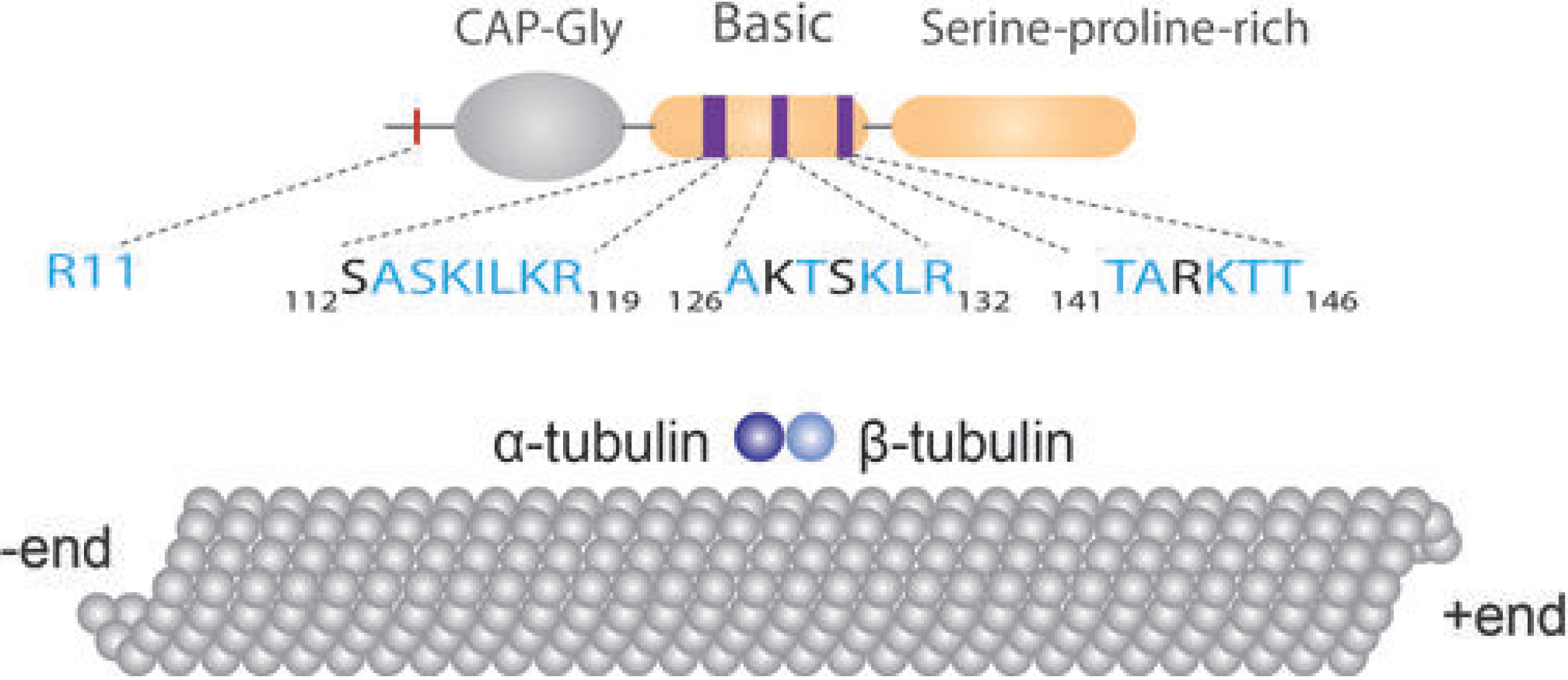
Schematic representation of p150^Glued^(1-191) and regions of its basic domain likely involved in binding to MTs. The segments comprised of residues S112–R119, A126–R132, and T141–R146 are structured and remain rigid upon binding with MTs. R11 is the only residue that is detected at 4 °C in the N-terminal segment 1-25.

## CONCLUSIONS

Our results demonstrate that the p150^Glued^(1-191) has stronger binding affinity with polymerized microtubules than the isolated CAP-Gly domain. Residues 108-191 of p150^Glued^(1-191) enhance the binding affinity and contain a second microtubule-binding region albeit it is largely unstructured in solution and dynamic upon binding to microtubules. Three short and rigid segments in the basic domain, S111–I116, A124–R132 and K144–T146 are predicted to form α-helical and β-sheet structures and are likely to encompass the microtubule binding site.

We conclude p150^Glued^(1-191) remains flexible upon binding to microtubules except for regions that are directly involved in the binding interactions. This conformational flexibility may be important for the p150^Glued^(1-191) interactions with microtubules. Our results lay the foundation for atomic-resolution structure characterization of dynactin’s p150^Glued^ subunit bound to polymerized microtubules.

## AUTHOR CONTRIBUTIONS

C.G., J.C.W. and T.P. designed the research. C.G. and T.P. performed experiments and analyzed data. C.G., J.C.W. and T.P. wrote the manuscript.

## ACKNOWLEDGEMENTS

We thank Dr. Si Yan for valuable discussions and initial work on the p150^Glued^(1-191) construct. We are grateful to Dr. Shi Bai for help in using solution NMR instruments and Shannon Modla at Bioimaging Center at the Delaware Institute of Technology for assistance with acquiring TEM images. We acknowledge the support of the NIH Grant P30GM110758 for the support to the core instrumentation infrastructure at the University of Delaware.

## Supporting Information

Tables of backbone resonance chemical shifts, chemical shift perturbations and ^31^P chemical shifts are listed in supplemental material. SDS-PAGE results for purification of dimers and cosedimentation assay of dimer and microtubules are included in Fig. S1.

